# What are we missing by using hydrophilic enrichment? Improving bacterial glycoproteome coverage using total proteome and FAIMS analysis

**DOI:** 10.1101/2020.07.22.216903

**Authors:** Ameera Raudah Ahmad Izaham, Ching-Seng Ang, Shuai Nie, Lauren E. Bird, Nicholas A. Williamson, Nichollas E. Scott

**Author notes:** **Corresponding Author** Dr Nichollas E. Scott, Address: Department of Microbiology and Immunology, University of Melbourne at the Peter Doherty Institute for Infection and Immunity, Melbourne 3000, Australia.

## Abstract

Hydrophilic Interaction Liquid Chromatography (HILIC) glycopeptide enrichment is an indispensable tool for the high-throughput characterisation of glycoproteomes. Despite its utility, HILIC enrichment is associated with a number of short comings including requiring large amounts of starting material, potentially introducing chemical artefacts such as formylation, and biasing/under-sampling specific classes of glycopeptides. Here we investigate HILIC enrichment independent approaches for the study of bacterial glycoproteomes. Using three Burkholderia species (*B. cenocepacia, B. dolosa* and *B. ubonensis*) we demonstrate that short aliphatic *O*-linked glycopeptides are typically absent from HILIC enrichments yet are readily identified in whole proteome samples. Using Field Asymmetric Waveform IMS (FAIMS) fractionation we show that at low compensation voltages (CVs) short aliphatic glycopeptides can be enriched from complex samples providing an alternative means to identify glycopeptides recalcitrant to hydrophilic based enrichment. Combining whole proteome and FAIMS analysis we show that the observable glycoproteome of these Burkholderia species is at least 30% larger than initially thought. Excitingly, the ability to enrich glycopeptides using FAIMS appears generally applicable, with the *N*-linked glycopeptides of *Campylobacter fetus subsp. fetus* also enrichable at low FAIMS CVs. Taken together, these results demonstrate that FAIMS provides an alternative means to access glycopeptides and is a valuable tool for glycoproteomic analysis.

## INTRODUCTION

Glycosylation is a ubiquitous protein modification found in all domains of life [1]. Over the last two decades significant progress has been made in our understanding of glycosylation targets leading to the establishment of the field of glycoproteomics [2, 3]. The growth of glycoproteomics has been driven by the increasing sensitivity and speed of mass spectrometry (MS) instrumentation as well as dramatic improvements in glycopeptide informatics [2–4]. These advancements now enable intact glycopeptide analysis – the study of glycopeptides decorated with their native [4], native-like [5, 6] or truncated [7, 8] glycoforms – and enables the qualitative and quantitative assessment of glycosylation at a site-specific resolution. Intact glycopeptide analysis has been critical for the study of glycosylation where the sites of modification are not predictable based on amino acid sequence such as mucin O-linked glycosylation [7, 8], *O*-mannosylation [9, 10] and *O*-glcNAcylation [11]. Although powerful, a key requirement for intact glycopeptide analysis is the ability to enrich/isolate glycopeptides for downstream analysis.

To date, MS based characterisation of intact glycopeptides has been heavily dependent on glycan centric glycopeptide enrichment methods [2–4, 12]. Enrichment has been proposed to be critical for concentrating glycopeptides in even simple protein digests [13] to overcome the suppression of glycopeptide ions by co-eluting non-glycosylated peptides [12]. A range of methods have been developed to enrich glycopeptides from mixtures [14] including lectin-affinity based approaches [15–17], reversible chemical coupling resins such as boronic acid based derivatives [18] and irreversible chemical strategies such as hydrazine based chemistry followed by carbohydrate selective hydrolysis [19]. One of the most widely used glycopeptide enrichment approaches is hydrophilic interaction liquid chromatography (HILIC) enrichment using zwitterionic hydrophilic interaction liquid chromatography (ZIC-HILIC) resins [20]. Enrichment using ZIC-HILIC is the result of hydrophilic partitioning of glycopeptides to an adsorbed “water layer” on the surface of the ZIC-HILIC resin [21, 22]. For glycopeptides, hydrophilic partitioning is thought to be driven largely by glycans making ZIC-HILIC enrichment compatible with a range of glycopeptides provided their glycans are sufficiently hydrophilic [23]. This enrichment approach has been used for nearly two decades [20, 24] for both eukaryotic [25–28] and prokaryotic [29–32] glycoproteomic studies. Although ZIC-HILIC enrichment is considered by some to be unbiased for eukaryotic *N*-linked glycoproteomics [12, 33], it has been previously shown to strongly favour large glycans [23] and show poor retention of *O*-linked glycopeptides [34]. This bias has important ramifications especially for the study of glycosylated substrates decorated with short glycans such as the glycoproteomes of Burkholderia species [31, 35].

Protein glycosylation is increasingly recognised as a highly conserved feature of bacterial physiology [36–39]. To date, studies of bacterial glycoproteomes have extensively used ZIC-HILIC enrichment [29–32, 40–43] to characterise glycosylation substrates and glycan diversity. Using ZIC-HILIC enrichment, we have previously shown over 100 proteins are glycosylated within *Burkholderia* species [44]. Despite this insight, the complete repertoire of most *Burkholderia* glycoproteomes are still unknown, hampering our understanding of the impacts observed when glycosylation is disrupted [31, 45, 46]. Within members of the Burkholderia genus, two linear trisaccharides composed of β-Gal-(1,3)-α-GalNAc-(1,3)-β-GalNAc and Suc-β-Gal-(1,3)-α-GalNAc-(1,3)-β-GalNAc (where Suc is Succinyl) are utilized for protein glycosylation [31, 35]. The short length of these two glycans raises notable concerns about potential bias in the observable glycoproteome within ZIC-HILIC enrichments [23, 34]. This concern, coupled with our recent identification of extensive formylation artefacts as a result of ZIC-HILIC enrichment [44], suggests alternative approaches – such as orthogonal fractionation – are required for the analysis of bacterial glycopeptides.

Ion Mobility Spectrometry (IMS) is an increasingly utilised gas-phase fractionation approach which enables the separation of analytes based on a combination of charge and collisional cross-section [47]. Within proteomics, Field Asymmetric Waveform IMS (FAIMS) is one of the most widely used IMS approaches [48] and has been used to increase proteome coverage [49–51], reduce ratio compression within isobaric labelling experiments [52], improve the localisation of PTMs [53, 54] as well as enrich modified peptides such as sumoylated [55] and cross-linked [56] peptides. Recently a new FAIMS interface was introduced for Orbitrap instruments: the FAIMS Pro interface [49–51]. This new implementation of FAIMS improves sensitivity and enables rapid compensation voltage (CV) switching, making it ideal for proteomics studies and gas-phase fractionation [49, 50]. By applying multiple CVs in a single experiment, referred to from here in as stepped FAIMS, this has been shown to increase protein identifications by up to 55% vs without FAIMS implementation [49–51]. Previous iterations of FAIMS have been shown to enable the separation of glycopeptide isomers [57, 58] as well as improve the mapping of bacterial glycosylation sites within single proteins [59]. Due to the large collisional cross-section of glycopeptides compared to unmodified peptides [60, 61], IMS has been suggested to provide a unique avenue for the sorting/enrichment of glycopeptides [60, 62]. However, to our knowledge FAIMS has not been previously used for glycoproteomics studies.

In this work we explore ZIC-HILIC independent approaches for the characterisation of bacterial glycoproteomes. By comparing the glycopeptides identified within three Burkholderia species (*B. cenocepacia, B. dolosa* and *B. ubonensis)* with and without ZIC-HILIC enrichment, we find that short aliphatic glycopeptides are under-represented within ZIC-HILIC enrichments. To improve the identification of these glycopeptides we explore the use of FAIMS and demonstrate that at low FAIMS CVs short/aliphatic glycopeptides can be readily enriched from total proteome samples. By combining multiple FAIMS CVs within a single analytical method and the analysis of total proteome samples, we demonstrate that the glycoproteome of these three Burkholderia species are at least 30% larger than previously thought. Similarly, glycopeptides decorated with large glycans such as the *N*-linked hexasaccharides of *Campylobacter fetus subsp. fetus* are equally enriched at low FAIMS CVs with the use of FAIMS providing comparable coverage of the glycoproteome using only a fraction of the starting material. The ability to access complementary glycoproteomes as well as enrich glycopeptides from low input amounts makes FAIMS a powerful tool for glycoproteomic analysis.

## EXPERIMENTAL PROCEDURES

### Bacterial strains and growth conditions

[63]. *B. cenocepacia* J2315; *B. dolosa* AU0158 and *B. ubonensis* MSMB22 strains were grown overnight on LB agar at 37°C as previously described [31]. *C. fetus subsp. fetus NCTC 10842* was grown on Brain-Heart Infusion medium (Hardy Diagnostics) with 5% defibrinated horse blood (Hemostat, Dixon, CA) under microaerobic conditions (10% CO_2_, 5% O_2_, 85% N_2_) at 37°C as previously reported [64]. Details on the strains, their origins, references and proteome databases are provided within Table 1.

**Table 1.** Strain list

| Strains | Source | Reference | Proteome database |
| --- | --- | --- | --- |
| *B. cenocepacia* LMG 16656 / J2315 | Human clinical specimen,<br>United Kingdom, 1989 | [88] | Uniprot database:<br>UP000001035 |
| *B. dolosa* AU0158 | Human clinical specimen,<br>USA unknown | [89] | Uniprot database:<br>UP000032886 |
| *B. ubonensis* MSMB22 | Soil isolate, Australia,<br>2001 | [89] | Burkholderia Genome<br>Database [90],<br>Strain number: 3404 |
| *C. fetus subsp. fetus* NCTC 10842 | Brain of sheep fetus,<br>France, 1952 | [91] | Uniprot database:<br>UP000001035 |

### Generation of bacterial lysates for glycoproteome analysis

Bacterial strains were grown to confluency before flooding plates with 5 mL of pre-chilled sterile phosphate-buffered saline (PBS) and bacterial cells collected by scraping. Cells were washed 3 times in PBS to remove media contaminates, then collected by centrifugation at 10,000 × *g* at 4°C for 10 min, and then snap frozen. Snap frozen cells were resuspended in 4% SDS, 100mM Tris pH 8.0, 20mM Dithiothreitol (DTT) and boiled at 95°C with shaking at 2000rpm for 10 min. Samples were clarified by centrifugation at 17,000 × *g* for 10 min, the supernatants were then collected, and protein concentrations determined by a bicinchoninic acid assay (Thermo Fisher Scientific, Waltham, MA, USA). 1mg of protein from each sample was acetone precipitated by mixing one volume of sample with 4 volumes of ice-cold acetone. Samples were precipitated overnight at −20°C and then spun down at 16,000 × *g* for 10 min at 0°C. The precipitated protein pellets were resuspended in 80% ice-cold acetone and precipitated for an additional 4 hours at −20°C. Samples were centrifuged at 17,000 × *g* for 10 min at 0°C, the supernatant discarded, and excess acetone driven off at 65°C for 5 min.

### Digestion of protein samples

Protein digestion was undertaken as previously described with minor alterations [29]. Briefly, dried protein pellets were resuspended in 6 M urea, 2 M thiourea in 40 mM NH_4_HCO_3_ then reduced for 1 hour with 20mM DTT, followed by alkylation with 40mM chloroacetamide for 1 hour in the dark. Samples were then digested with Lys-C (Wako Chemicals, Japan; 1/200 w/w) for 3 hours before being diluted with 5 volumes of 40 mM NH_4_HCO_3_ and digested overnight with sequencing grade modified trypsin (Promega, Madison, WI, USA; 1/50 w/w). Digested samples were acidified to a final concentration of 0.5% formic acid and desalted with 50 mg tC18 Sep-Pak columns (Waters corporation, Milford, MA, USA) according to the manufacturer’s instructions. tC18 Sep-Pak columns were conditioned with 10 bed volumes of Buffer B (0.1% formic acid, 80% acetonitrile), then equilibrated with 10 bed volumes of Buffer A* (0.1% TFA, 2% acetonitrile) before use. Samples were loaded on to equilibrated columns and then columns washed with at least 10 bed volumes of Buffer A* before bound peptides were eluted with Buffer B. Eluted peptides were aliquoted into samples for ZIC-HILIC enrichment or total proteome analysis, then dried by vacuum centrifugation and stored at −20°C.

### Reverse phase LC-MS/MS

Proteome samples were re-suspended in Buffer A* and separated using a two-column chromatography set up composed of a PepMap100 C18 20 mm × 75 μm trap and a PepMap C18 500 mm × 75 μm analytical column (Thermo Fisher Scientific). Samples were concentrated onto the trap column at 5 μL/min for 5 minutes with Buffer A (0.1% formic acid, 2% DMSO) and then infused into an Orbitrap Fusion™ Lumos™ Tribrid™ Mass Spectrometer (Thermo Fisher Scientific) at 300 nL/minute via the analytical column using a Dionex Ultimate 3000 UPLC (Thermo Fisher Scientific). For stepped CV and whole proteome experiments, 185-minute analytical runs were undertaken by altering the buffer composition from 2% Buffer B (0.1% formic acid, 77.9% acetonitrile, 2% DMSO) to 28% B over 150 minutes, then from 28% B to 40% B over 10 minutes, then from 40% B to 100% B over 2 minutes. The composition was held at 100% B for 3 minutes, and then dropped to 2% B over 5 minutes before being held at 2% B for another 15 minutes. These conditions are identical to the published ZIC-HILIC glycopeptide datasets [44] to enable the direct comparison of glycopeptide coverage. For static FAIMS CVs and associated no FAIMS control experiments, 125-minute analytical runs were undertaken by altering the buffer composition from 2% Buffer B (0.1% formic acid, 77.9% acetonitrile, 2% DMSO) to 28% B over 90 minutes, then from 28% B to 40% B over 10 minutes, then from 40% B to 100% B over 2 minutes. The composition was held at 100% B for 3 minutes, and then dropped to 2% B over 5 minutes before being held at 2% B for another 15 minutes. For all experiments, the Lumos™ Mass Spectrometer was operated in a data-dependent mode, automatically switching between the acquisition of a Orbitrap MS scan (120,000 resolution) every 3 seconds and Orbitrap MS/MS HCD scans of precursors (NCE 30%, maximal injection time of 60 ms, with an AGC of 200% and a resolution of 15,000). Glycan fragment ions (204.087; 366.1396 and 138.0545 m/z) product-dependent MS/MS analysis [65] was used to trigger three additional scans of potential glycopeptides; a Orbitrap EThcD scan (NCE 15%, maximal injection time of 250 ms with an AGC of 500% and a resolution of 30,000 using the extended mass range setting to improve the detection of high mass glycopeptide fragment ions [61]); an ion trap CID scan (NCE 35%, maximal injection time of 40 ms with an AGC of 200%) and a stepped collision energy HCD scan (using NCE 30%, 35%, 45% with a maximal injection time of 250 ms with an AGC of 500% and a resolution of 30,000). For FAIMS experiments, static FAIMS analysis was undertaken using CVs of −20; −30; −40; −50; −60; −70; −80 and −90 while stepped FAIMS experiments were undertaken switching between the CVs −25; −35 and −45. All experiments were collected in triplicate.

### Data Analysis - Glycopeptide identification

Raw data files were batch processed using Byonic v3.5.3 (Protein Metrics Inc. [66]) with the proteome databases denoted within Table 1. Data was searched on a desktop with two 3.00GHz Intel Xeon Gold 6148 processors, a 2TB SDD and 128 GB of RAM using a maximum of 16 cores for a given search. For all searches, a semi-tryptic N-ragged specificity was set, and a maximum of two missed cleavage events allowed. Carbamidomethyl was set as a fixed modification of cystine while oxidation of methionine was included as a variable modification. A maximum mass precursor tolerance of 5 ppm was allowed while a mass tolerance of up to 10 ppm was set for HCD fragments and 20 ppm for EThcD fragments. *Burkholderia* species searches were conducted allowing two *O*-linked glycans: Hex-HexNAc-HexNAc (elemental composition: C_22_O_15_H_36_N_2_, mass: 568.2115) and Suc-Hex-HexNAc-HexNAc (elemental composition: C_26_O_18_H_40_N_2_, mass: 668.2276), while *C. fetus fetus* searches were conducted allowing two *N*-linked glycans: Hex-HexNAc_4_-diNAcBac (elemental composition: C_48_O_29_H_78_N_6_, mass: 1202.4813) and HexNAc_5_-diNAcBac (elemental composition: C_50_O_29_H_81_N_7_, mass: 1243.5078) where diNAcBac is the bacterial specific sugar 2,4-diacetamido-2,4,6 trideoxyglucopyranose [64]. To ensure high data quality, technical replicates were combined using R (https://www.r-project.org/) and only glycopeptides with a Byonic score >300 were used for further analysis. This cut-off is in line with previous reports highlighting that score thresholds greater than at least 150 are required for robust glycopeptide assignments with Byonic [28, 67]. It should be noted that a score threshold of above 300 resulted in false discovery rates of less than 1% for all combined datasets.

### Data Analysis - iBAQ analysis

To assess the relative abundance of proteins, iBAQ analysis was undertaken. Unenriched proteome datasets were processed using MaxQuant v1.6.3.4 [68] searching against the reference *B. cenocepacia* strain J2315. Carbamidomethylation of cysteine was set as a fixed modification while oxidation of methionine and acetylation of protein N-termini were allowed as variable modifications. An enzyme specificity of “trypsin/P” was set with a maximum of 2 missed cleavage events. The resulting outputs were processed in the Perseus v1.5.0.9 [69] analysis environment to remove reverse matches and common protein contaminates prior to further analysis.

### Peptide analysis, data visualization and data sharing

The previously published ZIC-HILIC glycopeptide datasets [44] were used to compare the differences in glycopeptides observed with and without ZIC-HILIC enrichment. It should be noted that the proteome digests within this work are the same digests used for ZIC-HILIC enrichments in [44]. Analysis and visualization was undertaken using R (https://www.r-project.org/) with the glycopeptide physicochemical properties determined using the Peptides package [70] and data visualized using ggplot2 [71] and ggpubr packages. The aliphatic index within the Peptides package is calculated by assessing the relative volume occupied by aliphatic side chains within a peptide [72]. All mass spectrometry proteomics data (Raw data files, Byonic search outputs and R Scripts) have been deposited into the PRIDE ProteomeXchange Consortium repository [73, 74] with the dataset identifier: PXD020442 and are accessible with the reviewer account details: Username: Password: aoATFQMQ.

## RESULTS

### Discrete glycopeptide populations are observed within whole cell digests compared to ZIC-HILIC enriched samples

Recently we noted glycopeptides could be readily identified without enrichment across a range of bacterial proteomes [23, 78]. Given this, we sought to compare the glycopeptides observed with and without enrichment for bacterial species focusing on members of the Burkholderia genus such as *B. cenocepacia* J2315. Within whole cell digests of *B. cenocepacia* J2315 we find that only ~2% of all high-confidence PSMs (Byonic™ scores >300) correspond to glycopeptides (Supplementary Figure 1A, Supplementary Table 1). Although a low proportion of all PSMs this still corresponds to hundreds of individual glycopeptide PSMs identified within analytical runs (Supplementary Figure 1B, Supplementary Table 1). Intuitively, we assumed that within these whole cell digests of *B. cenocepacia* J2315, the observed glycopeptides would correspond to previously identified glycopeptides within ZIC-HILIC enrichments, yet this was not the case. Surprisingly, the majority of glycopeptides identified within whole cell digests corresponded to novel glycoproteins/glycopeptides (Figure 1A and B), with manual inspection of previously unreported glycopeptides supporting the correctness of these assignments (Supplementary Figure 2A-H). These results demonstrate that different subsets of the *B. cenocepacia* J2315 glycoproteome are observed with and without ZIC-HILIC enrichment.

**Figure 1:**
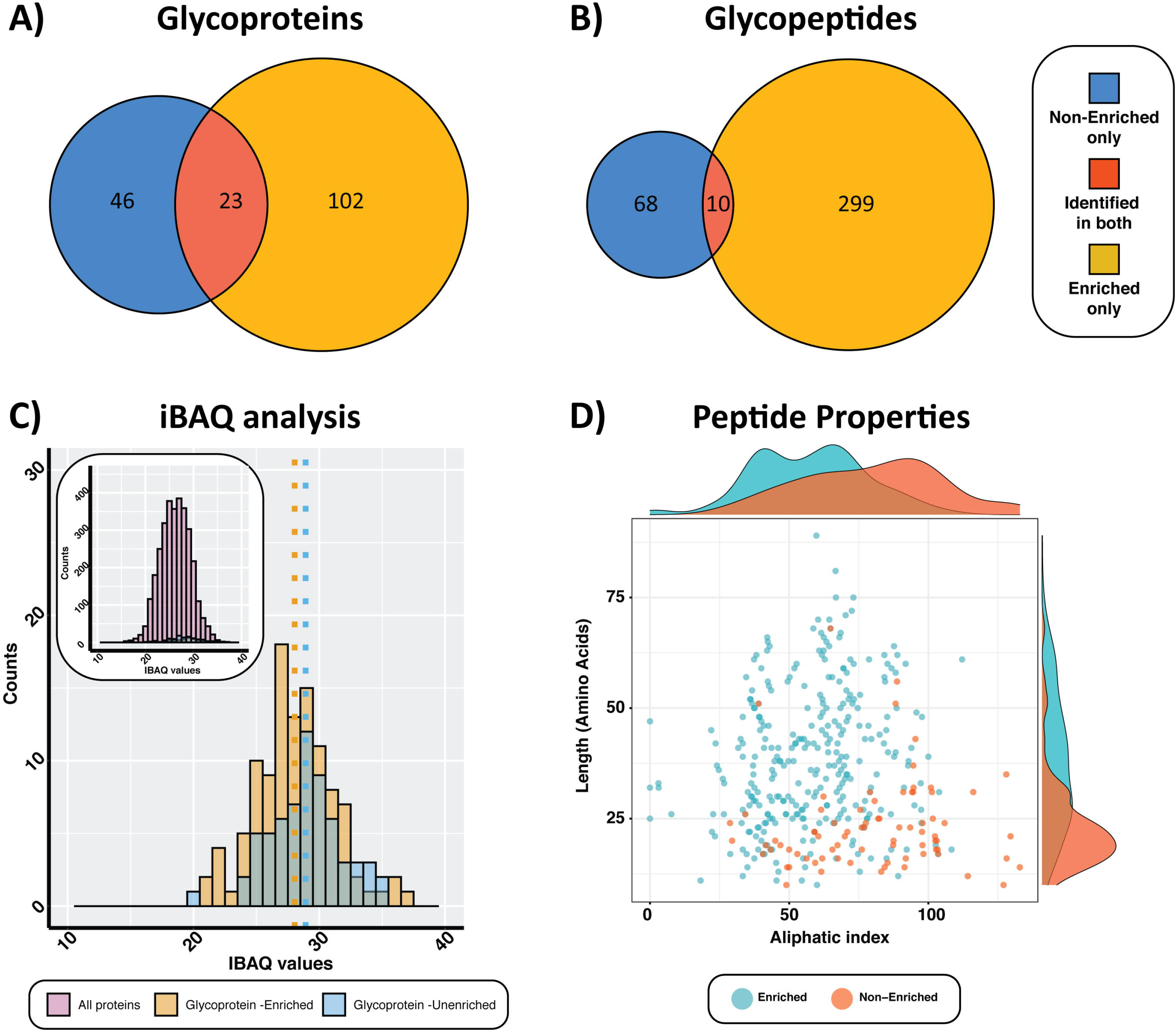
Discrete glycopeptides are observed within unenriched and ZIC-HILIC enriched bacterial samples. **A and B)** The overlap of glycoproteins (A) and glycopeptides (B) identified without and with ZIC-HILIC enrichment demonstrates the majority of glycopeptides/glycoproteins observed within unenriched samples are unique to those observed within ZIC-HILIC enrichments. **C)** iBAQ measurements demonstrate only modest differences in the relative abundance of glycoproteins observed with and without ZIC-HILIC enrichment. The mean iBAQ values for each group are denoted by the dotted lines. **D)** The physiochemical properties of glycopeptides observed with and without ZIC-HILIC enrichment demonstrates glycopeptides observed without enrichment are shorter and more aliphatic then glycopeptides observed with ZIC-HILIC enrichment.

To understand the differences in the glycopeptides identified with and without ZIC-HILIC enrichment we first compared the relative abundances of glycoproteins using iBAQ values derived from whole cell digests (Figure 1C, Supplementary Table 2). We reasoned that the dramatic difference in the glycopeptide profiles may be linked to glycoprotein abundances with ZIC-HILIC enrichment potentially accessing glycopeptides from lower abundance glycoproteins. Although the iBAQ values of glycoproteins observed within enriched samples are slightly lower (*P* = 0.0036; mean iBAQ value of glycoproteins identified in enriched samples = 28.045 compared to the mean iBAQ value of glycoproteins identified within unenriched samples = 28.918, Supplementary Table 3), the difference in the abundance is modest (Figure 1C), suggesting that the difference in the *B. cenocepacia* glycopeptides profiles observed between samples is independent of protein abundance. As previous reports have suggested that peptide sequences can influence ZIC-HILIC glycopeptide enrichment [23, 75] we examined the physiochemical properties of the peptide sequences revealing marked differences in the glycopeptides observed between samples (Figure 1D). We found that glycopeptides observed within whole cell digests are both more aliphatic and typically shorter in length, while the majority of ZIC-HILIC enriched glycopeptides are both longer and less aliphatic (*P =* 5.10*10^15^ and 6.68*10^15^ aliphaticity and length respectively, Supplementary Table 4). Combined these results support that differences in the physiochemical properties of glycopeptides influences their ability to be enriched by ZIC-HILIC enrichment.

### ZIC-HILIC incompatible glycopeptides are readily accessible using FAIMS fractionation

The findings that ZIC-HILIC enrichment provides access to only a subset of Burkholderia glycopeptides suggests that alternative approaches are required for comprehensive analysis of Burkholderia glycoproteomes. As glycopeptides were readily identified within whole cell digests of *B. cenocepacia*, we reasoned that an orthogonal fractionation approach may identify previously overlooked glycopeptides. Since FAIMS enables gas-phase fractionation based on charge/collision cross section [49, 50], we examined the charge state distributions of glycopeptides identified from whole cell digests compared to ZIC-HILIC enrichments (Figure 2A and B). We noted that glycopeptides observed within the whole cell digest were predominantly of low charge states (<+3). As previous work [50] has shown that +3 and +2 peptides are identified across CVs of −20 to −120 with apexes at −80 and −45 respectively this suggested low FAIMS CVs would enable enrichment of potential low charge state glycopeptides. To assess this, we assayed the ability of *B. cenocepacia* glycopeptides to be identified under differing FAIMS CVs from −20 to −90 (Figure 2C, Supplementary Table 5 and 6). Consistent with our hypothesis we found that FAIMS CVs of less then −50 readily enabled the isolation of glycopeptides (Figure 2C, Supplementary Figure 3A). Interestingly, we also found that within individual analytical runs (denoted with dots in Figure 2C) with FAIMS CVs of −20, −30 and −40 more unique glycopeptides were identified per run than analysis without FAIMS. However, when the results of triplicate runs were combined (denoted by the bars in Figure 2C), only the FAIMS CV −30 outperformed analysis without FAIMS (Figure 2C). Similar to whole proteome digests, FAIMS-fractionated glycopeptides typically corresponded to novel glycoproteins/glycopeptides that were not captured by ZIC-HILIC enrichments (Figure 2D and E) and were again shorter and more aliphatic then ZIC-HILIC enriched glycopeptides (Supplementary Figure 3B). Within FAIMS CVs we noted that, consistent with previous reports [50], the overlap between 10 CV fractions was modest with a large proportion of glycopeptides unique to a single CV (Figure 3A and B). Taken together these results suggest that low FAIMS CVs provide access to ZIC-HILIC inaccessible *B. cenocepacia* glycopeptides yet the coverage of the glycoproteome within a single FAIMS CV is limited.

**Figure 2.**
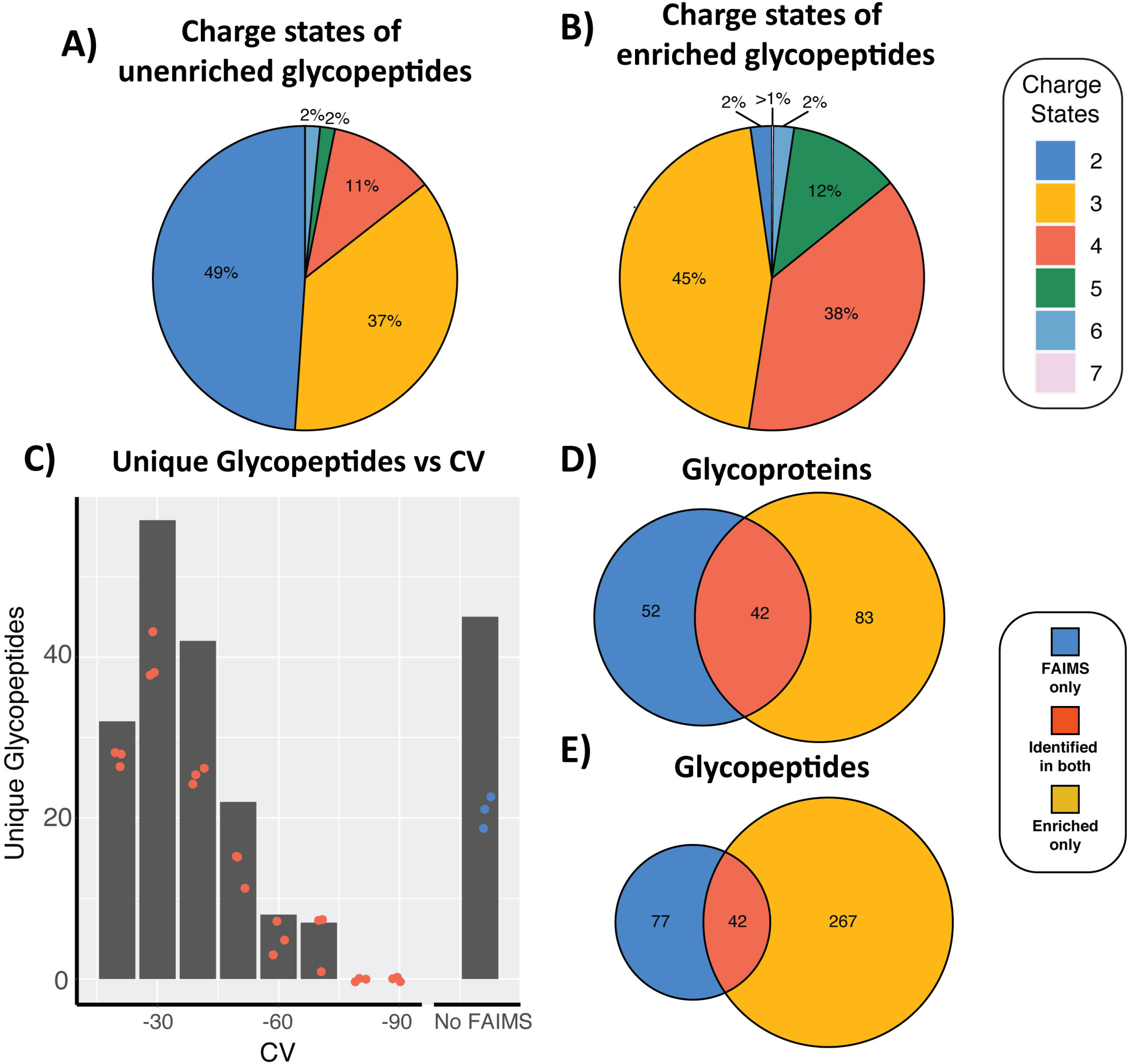
FAIMS fractionation enables the enrichment of glycopeptides within *B. cenocepacia J2315*. **A and B)** Charge state distributions of glycopeptides observed without ZIC-HILIC enrichment (A) and glycopeptides observed with ZIC-HILIC enrichment (B). **C)** 2 hr analytical gradients of FAIMS fractionation at different CVs demonstrate glycopeptides are enriched at low FAIMS CVs compared to 2 hr analytical gradients of analysis without FAIMS fractionation. Red dots denote the numbers of unique glycopeptides in individual replicates at each FAIMS CV. Blue dots denote the numbers of unique glycopeptides in individual replicates from analysis without FAIMS. Bars represent the total number of unique glycopeptides identified under each condition. **D and E)** FAIMS fractionated samples enable the identification of glycoproteins (D) and glycopeptides (E) not observed within ZIC-HILIC enrichments.

**Figure 3.**
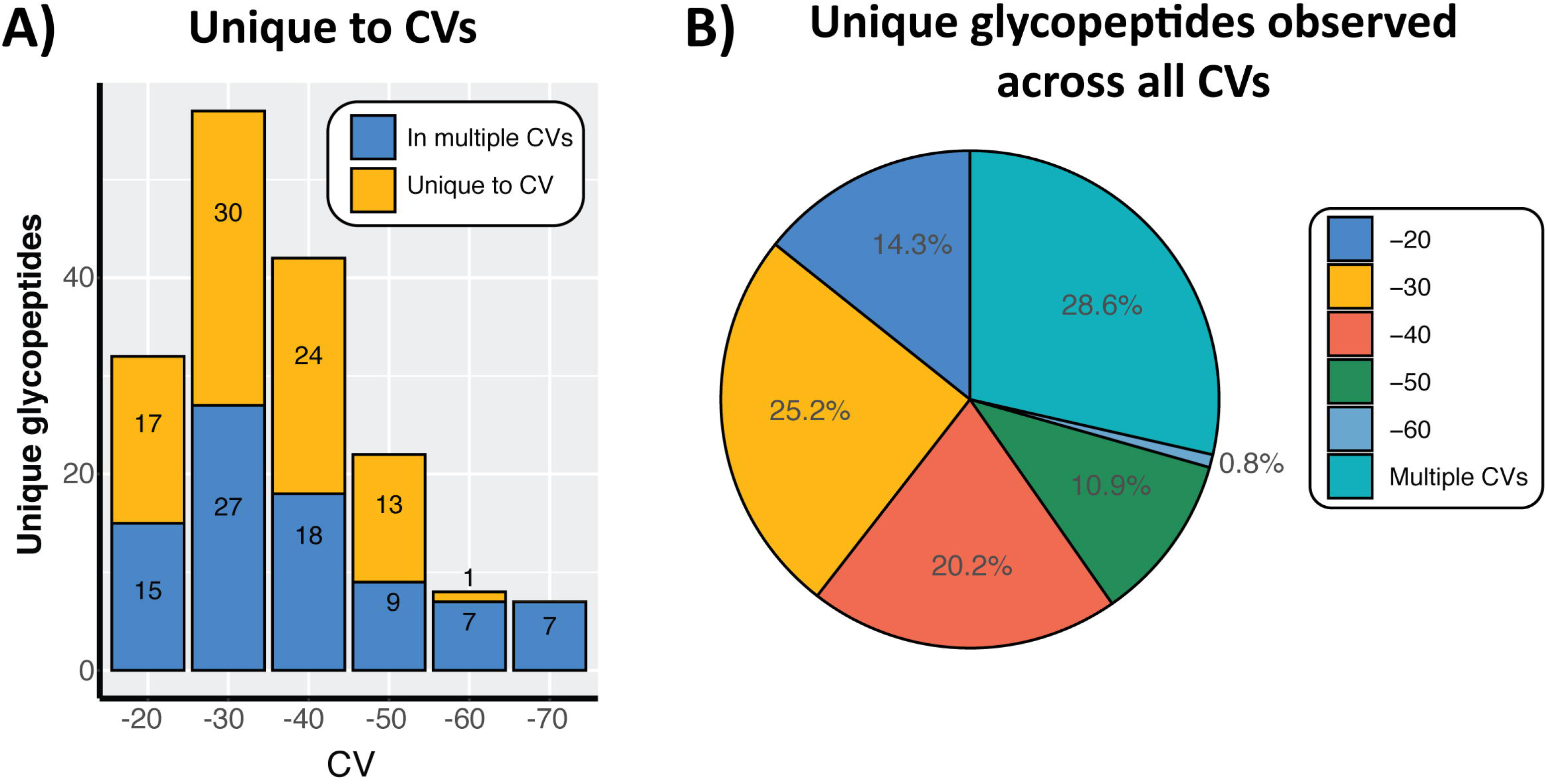
FAIMS CV fractions provide access to unique glycopeptide pools. **A)** Analysis of static CVs observed to contain glycopeptides demonstrates that a large proportion of glycopeptides are unique to a single CV fraction. **B)** Pie chart of unique glycopeptides observed within static FAIMS CV fractions. The majority of all glycopeptides are observed in CVs below −50.

### Using stepped FAIMS improves the glycoproteome coverage of Burkholderia species

Since individual FAIMS CVs provided access to only a subset of *B. cenocepacia* glycopeptides, we reasoned that using stepped FAIMS may improve glycopeptide identifications. Comparing the performance of stepped FAIMS, unenriched and ZIC-HILIC enriched digests we find that stepped FAIMS leads to a >30% increase in unique glycopeptides compared to unenriched samples (Figure 4A, Supplementary Table 7). As with static FAIMS CVs this improvement was predominantly within glycopeptides not previously observed within ZIC-HILIC enrichments with >60% of glycopeptides unique to stepped FAIMS (Figure 4B). Interestingly, we find that the glycopeptides identified in both stepped FAIMS and unenriched samples provide access to unique glycopeptide subsets compared to ZIC-HILIC enrichments (Figure 4C). Analysis of the physiochemical properties of glycopeptides observed using each approach highlights that the glycopeptides observed within *B. cenocepacia* with stepped FAIMS and unenriched samples are largely similar, favouring shorter and more aliphatic glycopeptides compared to the larger and less aliphatic glycopeptides observed with ZIC-HILIC enrichment (Figure 4D, Supplementary Table 8). The modest overlap of unique glycopeptides observed with stepped FAIMS and analysis without FAIMS fractionation illustrates that the glycoproteome of *B. cenocepacia* is larger than initially thought [31, 44], and that ZIC-HILIC enrichment fails to capture a significant proportion of the Burkholderia glycoproteome.

**Figure 4.**
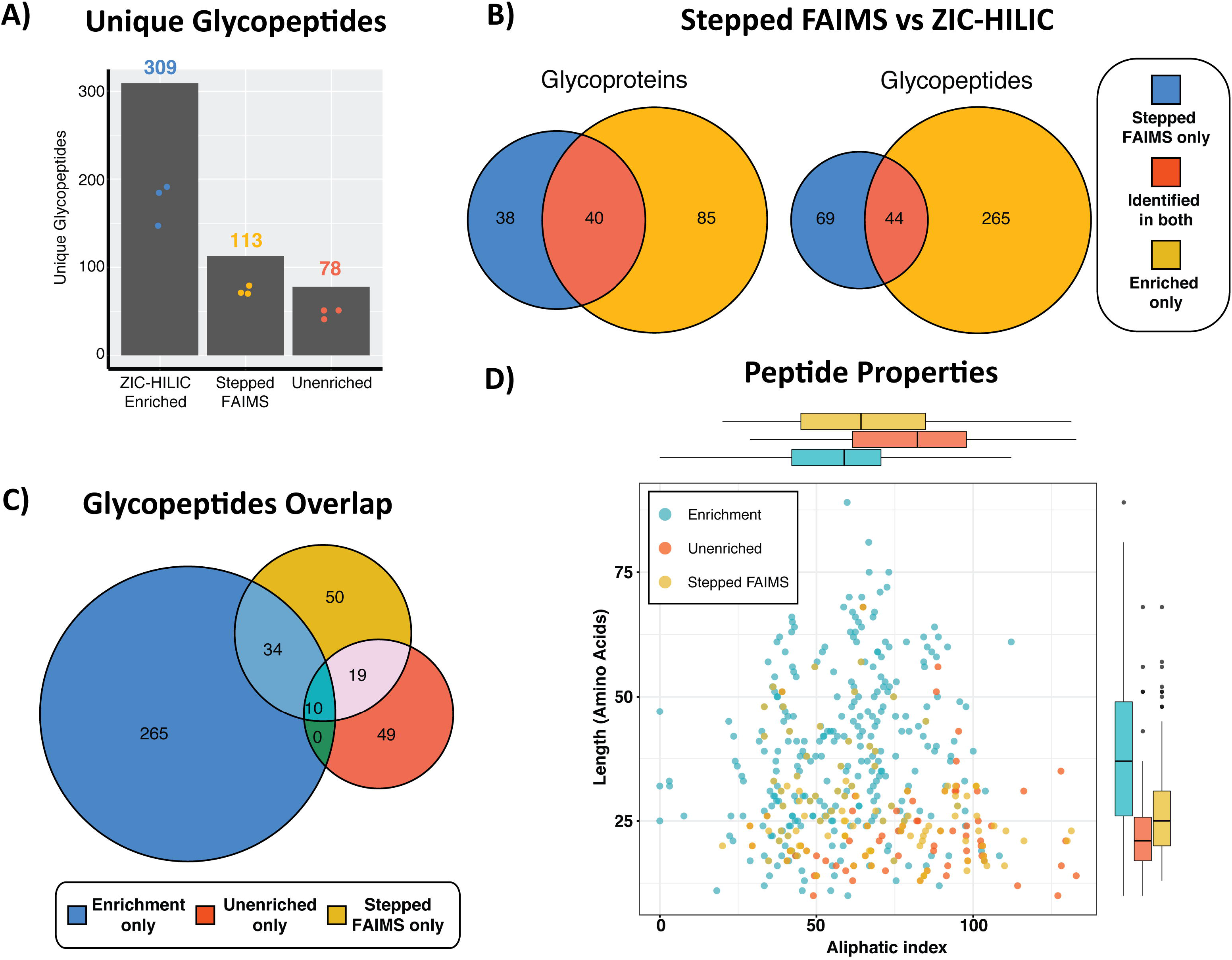
Stepped FAIMS Improves the identification of glycopeptides within *B. cenocepacia.* **A)** Comparison of 3 hr analytical gradients of ZIC-HILIC enrichments, stepped FAIMS and unenriched digests. Stepped FAIMS improves the identification of unique glycopeptides compared to unenriched samples. Individual replicates are denoted by dots and the combined number of unique glycopeptides within an approach are denoted by bars **B)** Comparison of the overlap in glycoproteins and glycopeptides identified with stepped FAIMS compared to ZIC-HILIC enrichments reveals stepped FAIMS enables access to glycopeptides/glycoproteins not observed within ZIC-HILIC enrichments. **C)** Overlap of glycopeptides observed within ZIC-HILIC enrichments, stepped FAIMS and analysis without FAIMS fractionation identifies that unique pools of glycopeptides are identified within each approach. **D)** The physiochemical properties of glycopeptides demonstrate that stepped FAIMS and analysis of unenriched digests allows the identification of typically more aliphatic and shorter glycopeptides.

To further confirm this bias within ZIC-HILIC enrichments we assessed two additional Burkholderia species recently assessed using ZIC-HILIC based enrichment: *B. dolosa and B. ubonensis* [44]. Consistent with our observations within *B. cenocepacia* both stepped FAIMS and unenriched samples revealed multiple unique glycopeptides/glycoproteins absent within ZIC-HILIC enrichments (Figure 5A to B; Supplementary Tables 9-13). As with *B. cenocepacia*, these newly observed glycopeptides in each Burkholderia species are typically shorter and/or more aliphatic than glycopeptides observed within ZIC-HILIC enrichments (Supplementary Figure 4). These results demonstrate that at least 30% of the glycoproteome of Burkholderia species are not observed within ZIC-HILIC enrichments, making orthogonal approaches such as stepped FAIMS valuable tools to access uncharacterised regions of Burkholderia glycoproteomes.

**Figure 5.**
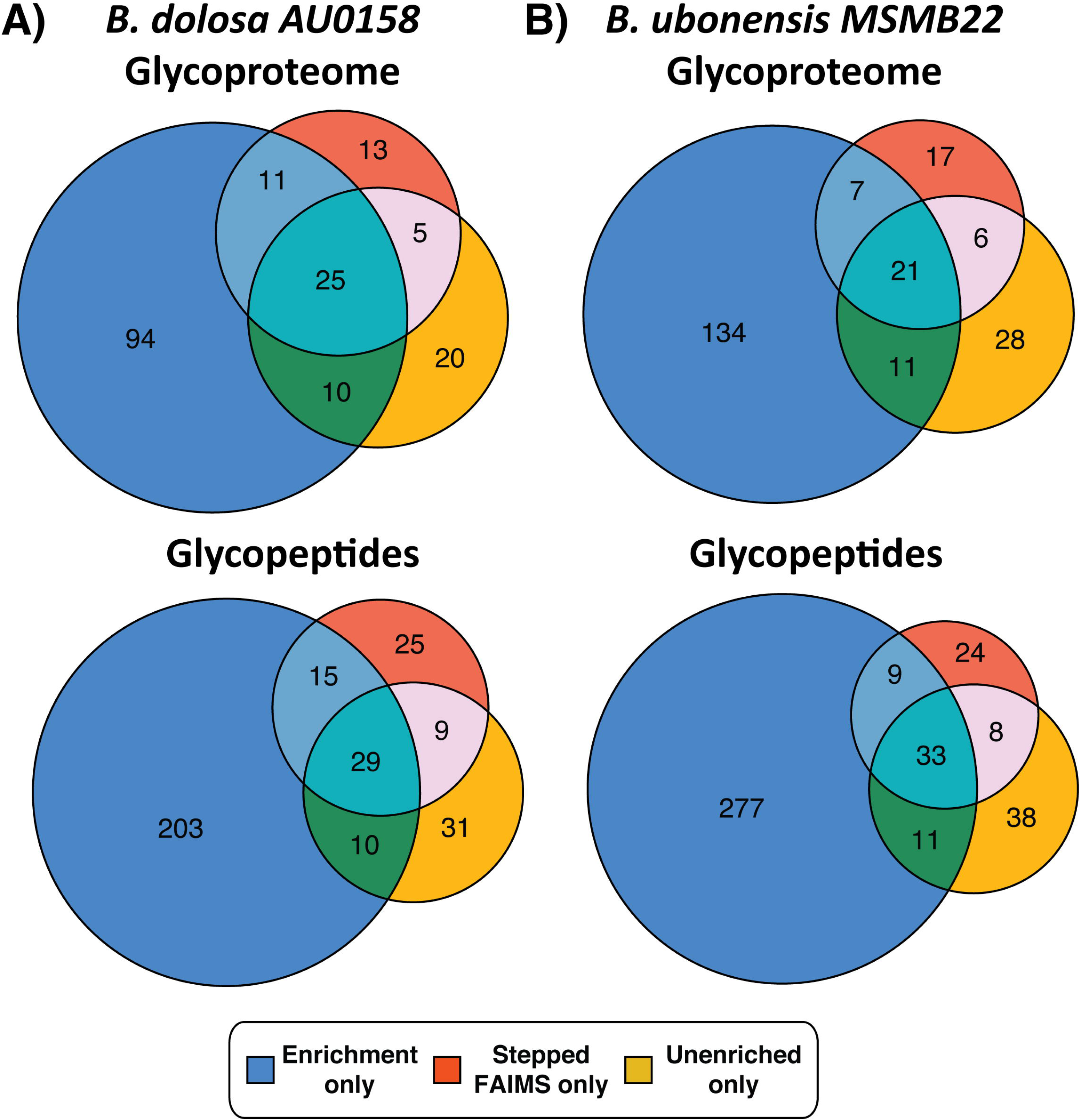
Comparison of glycoproteome coverage of Burkholderia species with ZIC-HILIC enrichment, Stepped FAIMS and analysis of unenriched samples. Analysis of *B. dolosa and B. ubonensis* demonstrate that multiple novel glycopeptides/glycoproteins are readily identified using stepped FAIMS and unenriched whole proteome samples.

### FAIMS fractionation improves the identification of bacterial *N*-linked glycoproteomes

Finally, as FAIMS improved the identification of glycopeptides within Burkholderia glycoproteomes, we wondered if other bacterial glycoproteomes would also benefit from FAIMS fractionation. To assess this we used *C. fetus subsp. fetus NCTC 10842* which is known to *N*-glycosylate proteins with two glycans composed of β-GlcNAc-α1,3-[GlcNAc1,6-]GlcNAc-α1,4-GalNAc-α1,4-GalNAc-α1,3-diNAcBac and β-GlcNAc-α1,3-[Glc1,6-]GlcNAc-α1,4-GalNAc-α1,4-GalNAc-α1,4-diNAcBac [64]. As with the trisaccharide decorated glycopeptides of Burkholderia, we noted a marked enrichment of *C. fetus subsp. fetus* glycopeptides within low CVs FAIMS fractions compared to matching analytical runs without FAIMS (Figure 6A, Supplementary Table 14 and 15). As observed within Burkholderia glycoproteomes, the use of stepped FAIMS improved the detection of unique glycopeptides compared to unenriched samples with matching analytical gradients (Figure 6B, Supplementary Table 16). Excitingly, we find that stepped FAIMS enabled the identification of a large proportion of previously identified glycoproteins and glycopeptides from ZIC-HILIC enrichments (Figure 6C, Supplementary Table 17) while requiring significantly less material (2μg per stepped FAIMS run compared to 300μg per ZIC-HILIC enrichment). These results demonstrate that large glycan-containing glycopeptides are enriched at low FAIMS CVs in a similar manner to glycopeptides decorated with shorter glycans, and this supports the general utilisation of FAIMS fractionation for isolating bacterial glycopeptides.

**Figure 6.**
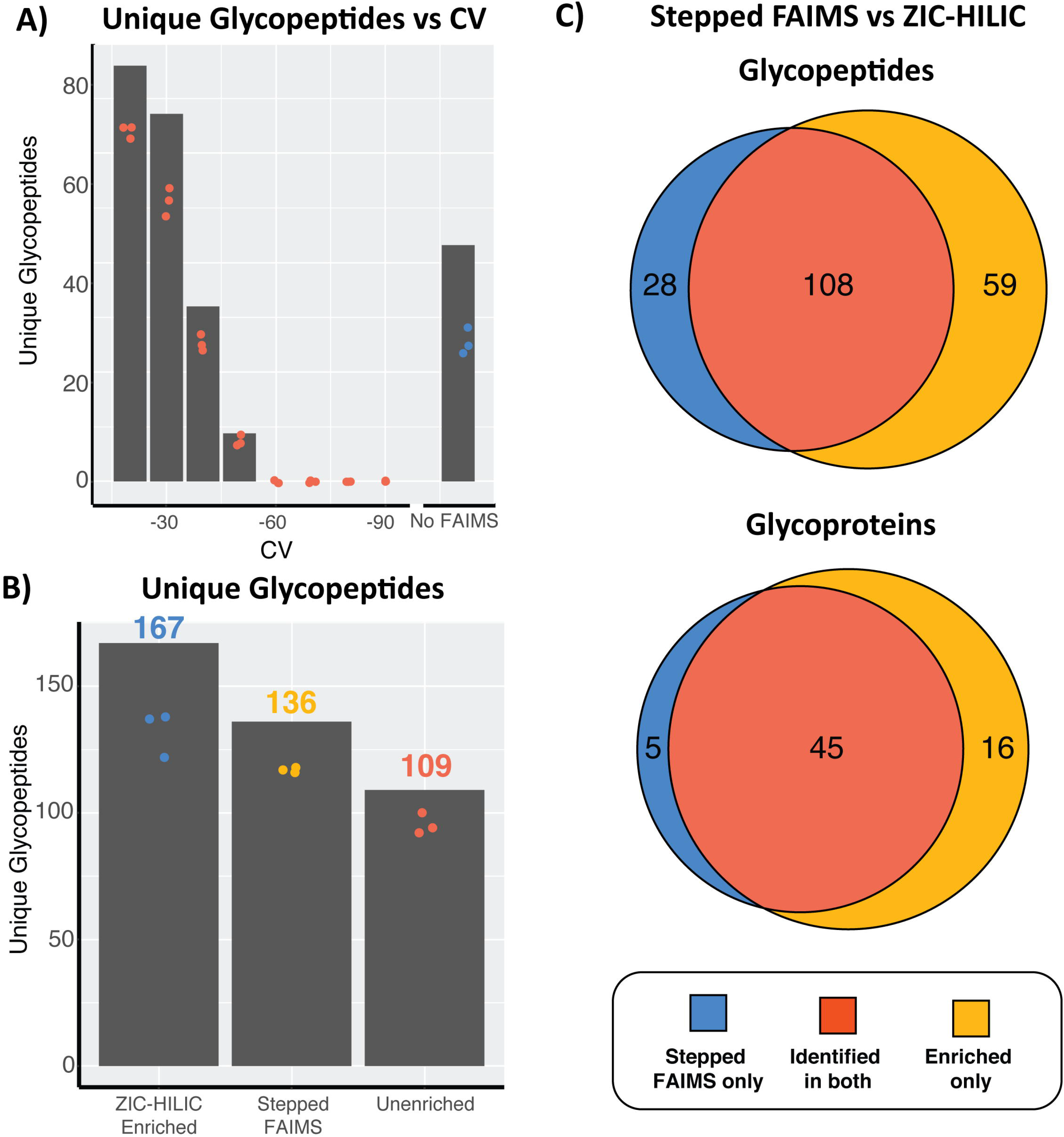
FAIMS fractionation of *C. fetus subsp. fetus N*-linked glycopeptides. **A)** Static FAIMS CVs demonstrate glycopeptides are enriched at low FAIMS CVs compared to analysis without FAIMS fractionation within 2hr analytical runs. Red dots denote the numbers of unique glycopeptides in individual replicates in each FAIMS CV. Blue dots denote the numbers of unique glycopeptides in individual replicates from analysis without FAIMS. Bars represent the total number of unique glycopeptides identified under each condition. **B)** Comparison of 3hr analytical runs of ZIC-HILIC enrichment, stepped FAIMS and analysis of unenriched whole proteome samples demonstrates an improvement in unique glycopeptides identified with stepped FAIMS compared to analysis without FAIMS fractionation. **C)** Comparison of the overlap in glycoproteins and glycopeptides identified with stepped FAIMS compared to ZIC-HILIC enrichments. At both the glycopeptide and glycoprotein level, stepped FAIMS enables identification of >60% of the known glycoproteome.

## Discussion

Over the last decade, glycopeptide-based enrichment approaches have increasingly been used to track and characterise microbial glycosylation events [42, 44, 64, 76]. Although enrichment strategies enable high-throughput analysis of glycosylation, we demonstrate here that hydrophilic enrichment provides only partial access to some microbial glycoproteomes. In light of these findings we evaluated the application of FAIMS fractionation using the FAIMS Pro interface [49–51] as an orthogonal approach for probing bacterial glycosylation. We find that for bacterial *N*- and *O*-linked glycans, low FAIMS CVs enables the enrichment of glycopeptides from whole proteome samples. By utilising a stepped FAIMS approach in concert with direct analysis of whole cell digests we show that the glycoproteomes of three Burkholderia species are >30% larger than initially thought and that the majority of the known glycoproteome of *C. fetus subsp. fetus* can be directly monitored from whole proteome digests.

To date, multiple studies have shown that glycosylation events can be readily detected within proteome samples using variable modification searches [77, 78] or open searching approaches [79–81]. These previous findings and our work here highlight that a wealth of glycoproteome information can be gained from unenriched samples, despite the known issue of glycopeptide ion suppression [12]. Experimentally, it has been shown that glycopeptide signals can be suppressed up to ten-fold compared to unmodified forms of the identical peptide [82] yet this issue is addressable. As current state of the art MS instruments enable the identification of potential glycopeptides by the presence of known sugar oxonium ions [65] this allows glycopeptide spectra to be re-collected with altered settings generating high-quality MS/MS data for even low abundance signals. Since FAIMS fractionation occurs post ionisation, it should be noted that ion suppression is not prevented but is likely being mitigated by the improvement in the signal to noise of analytes. As the amount of ion suppression is impacted by the complexity of the attached glycans of glycopeptides [82], it is reasonable to assume that some glycoproteomes will be more amenable to FAIMS analysis then others. Our work here demonstrates that glycopeptides of Burkholderia species and *C. fetus subsp. fetus* appear readily amenable to FAIMS enrichment, although further work will be required to establish general trends for specific classes of glycosylation.

The ability to enrich glycopeptides independent of off-line glycan-centric enrichment strategies opens up exciting possibilities for the study of previously recalcitrant glycoproteomes. Previous comparative genomic [83] as well as *ex vivo* studies [84] have identified a number of potential bacterial glycosylation systems for which the native substrates and glycans have yet to be defined. For example, in *Vibrio cholerae* a functional *O*-linked glycosylation system has been identified [84] and shown to be required for cholera toxin secretion [85], yet the glycoproteins of this system are still unknown. Due to the diversity of glycans used for bacterial protein glycosylation [29, 30, 35, 41, 42, 59, 64, 76, 86, 87] it is not surprising that a single approach such as ZIC-HILIC enrichment is inapplicable to all glycosylation systems or substrates within a single glycoproteome. Indeed, within this study we note that the bias within Burkholderia ZIC-HILIC enrichments does not appear to occur within ZIC-HILIC enrichments of *C. fetus subsp. fetus* (Supplementary Figure 5, Supplementary Table 17). This difference is likely due to the larger more hydrophilic glycans of *C. fetus subsp. fetus* which are retained more effectively during ZIC-HILIC enrichment. Thus, the diversity within microbial glycosylation systems means that multiple approaches are likely needed, with FAIMS providing an orthogonal approach for undertaking high throughput microbial glycoproteomic analysis.

The observation that FAIMS enables the enrichment of glycopeptides at a proteome scale has not explicitly been shown before although multiple IMS studies have highlighted this potential. In 2013 the Hill group demonstrated the ability of IMS to separate glycosylated and non-glycosylated components based on differences in collisional cross section [62], noting that this could provide a novel strategy for high throughput glycopeptide identification. This concept has been echoed in more recent studies which have shown that the ion mobility of low charge state glycopeptides can be dramatically impacted by the presence of glycans [60]. Our finding that FAIMS enables access to novel glycopeptides is consistent with previous studies of single glycoproteins [59] and we demonstrate that this can be extended to the analysis of glycoproteomes. A clear benefit of the recent FAIMS Pro interface is the rapid CV switching enabling multiple CVs to be accessible within a single analytical framework. We find stepped FAIMS approaches improves the detection of unique glycopeptides (Figure 4A and 6A) yet it should be noted that the optimal stepping regimes for glycopeptide identification is still unclear.

It is plausible that further improvements in glycoproteome coverage may be possible by optimising the CV switching to ensure minimal overlap between steps, while still focusing on the low CV region where glycopeptides are enriched. Our results provide support for the notion that FAIMS can improve our ability to study previously overlooked regions of glycoproteomes, warranting further investigation of these methods for the field of glycoproteomics.

## ASSOCIATED CONTENT

### Supporting Information

**Supplementary document:** Containing Supplementary figures 1 to 5

**Supplementary Figure 1: Identification of glycopeptides within** *B. cenocepacia* **J2315 whole proteome digests.** A) Pie charts showing all unique PSMs identified with scores above 300 B) Number of glycopeptides PSMs (Byonic Score >300) identified within each unenriched sample.

**Supplementary Figure 2: Manual assessment of glycopeptides identified with** *B. cenocepacia* **J2315 whole proteome digests.** HCD and ETD fragmentation of novel glycoproteins A-B) B4E7L5; C-D) B4EBZ3; E-F) B4E732 and G-H) B4EB92.

**Supplementary Figure 3: Analysis of** *B. cenocepacia* **glycopeptide identified within static FAIMS fractions.** A) Scatter plot of CV vs m/z of glycopeptide identified. B) Peptide properties of glycopeptide identified across static FAIMS CVs vs HILIC enriched glycopeptides

**Supplementary Figure 4: Comparisons of glycopeptide length vs aliphaticity across the Burkholderia species** *B. dolosa and B. ubonensis*. Within strains examined difference in aliphaticity properties or length are observed from glycopeptides identified within stepped FAIMS and unenriched samples compared to ZIC-HILIC enrichments.

**Supplementary Figure 5: Comparisons of glycopeptide length vs aliphaticity within** *C. fetus subsp. fetus glycopeptide*. No differences in the aliphatic properties or length are observed from glycopeptides identified within stepped FAIMS and unenriched samples compared to ZIC-HILIC enrichments.

**Supplementary Tables:** All Supplementary tables are provided within a single zip file

**Supplementary table 1: Combined Byonic searches of** *B. cenocepacia* **J2315 whole proteome digests.** The combined Byonic results of three technical replicates of *B. cenocepacia* J2315 showing all glycopeptides identified with a Byonic score over 300.

**Supplementary table 2: Maxquant analysis of** *B. cenocepacia* **J2315**. Maxquant analysis of three technical replicates of *B. cenocepacia* J2315 whole proteome digests.

**Supplementary table 3: iBAQ measurements of** *B. cenocepacia* **J2315**. iBAQ values for all glycoproteins identified within enriched and unenriched samples of *B. cenocepacia* J2315 whole proteome digests.

**Supplementary table 4: Peptide properties of** *B. cenocepacia* **J2315 glycopeptides identified within and without ZIC-HILIC enrichments**. For each glycopeptide identified within unenriched and ZIC-HILIC enriched samples the length and aliphatic index is provided.

**Supplementary table 5: Combined Byonic searches of** *B. cenocepacia* **J2315 without FAIMS**. The combined Byonic results of three technical replicates of *B. cenocepacia* J2315 showing all glycopeptides identified with a Byonic score over 300.

**Supplementary table 6: Combined Byonic searches of** *B. cenocepacia* **J2315 with static FAIMS CVs**. The combined Byonic results of each CV from −20 to −90 (three technical replicates of each) for *B. cenocepacia* J2315 showing all glycopeptides identified with a Byonic score over 300.

**Supplementary table 7: Combined Byonic searches of** *B. cenocepacia* **J2315 with stepped FAIMS**. The combined Byonic results for three technical replicates of *B. cenocepacia* J2315 with stepped FAIMS showing all glycopeptides identified with a Byonic score over 300.

**Supplementary table 8: Peptide properties of glycopeptides identified within** *B. cenocepacia* **J2315 across analysis approaches**. For each glycopeptide identified within stepped FAIMS, unenriched and ZIC-HILIC enriched samples the length and aliphatic index are provided.

**Supplementary table 9: Combined Byonic searches of** *B. ubonensis MSMB22* **with stepped FAIMS**. The combined Byonic results for three technical replicates of *B. ubonensis MSMB22* with stepped FAIMS showing all glycopeptides identified with a Byonic score over 300.

**Supplementary table 10: Combined Byonic searches of** *B. ubonensis MSMB22* **without FAIMS**. The combined Byonic results of three technical replicates of *B. ubonensis MSMB22* showing all glycopeptides identified with a Byonic score over 300.

**Supplementary table 11: Combined Byonic searches of** *B. dolosa AU0158* **with stepped FAIMS**. The combined Byonic results for three technical replicates of *B. dolosa AU0158*with stepped FAIMS showing all glycopeptides identified with a Byonic score over 300.

**Supplementary table 12: Combined Byonic searches of** *B. dolosa AU0158* **without FAIMS**. The combined Byonic results of three technical replicates of *B. dolosa AU0158* showing all glycopeptides identified with a Byonic score over 300.

**Supplementary table 13: Peptide properties of glycopeptides identified within** *B. ubonensis MSMB22* **and** *B. dolosa AU0158* **across analysis approaches**. For each glycopeptide identified within stepped FAIMS, unenriched and ZIC-HILIC enriched samples the length and Aliphatic index is provided.

**Supplementary table 14: Combined Byonic searches of** *C. fetus subsp. fetus NCTC 10842* **with static FAIMS CVs**. The combined Byonic results of each CV from −20 to −90 (three technical replicates) for *C. fetus subsp. fetus NCTC 10842* showing all glycopeptides identified with a Byonic score over 300.

**Supplementary table 15: Combined Byonic searches of** *C. fetus subsp. fetus NCTC 10842* **without FAIMS**. The combined Byonic results of three technical replicates of *C. fetus subsp. fetus NCTC 10842* showing all glycopeptides identified with a Byonic score over 300.

**Supplementary table 16: Combined Byonic searches of** *C. fetus subsp. fetus NCTC 10842* **with stepped FAIMS**. The combined Byonic results for three technical replicates of *C. fetus subsp. fetus NCTC 10842* with stepped FAIMS showing all glycopeptides identified with a Byonic score over 300.

**Supplementary table 17: Peptide properties of glycopeptides identified within** *C. fetus subsp. fetus NCTC 10842* **across analysis approaches**. For each glycopeptide identified within stepped FAIMS, unenriched and ZIC-HILIC enriched samples the length and Aliphatic index is provided.

## AUTHOR INFORMATION

### Author contributions

A.R.A.I prepared all sample for MS analysis. C.S.A, S.N and N.A.W maintained MS instrumentation and provided access to critical resources. L.E.B assisted in manuscript preparation. N.E.S conceived, designed all experiments, performed analysis, and undertook the preparation of the manuscript.

### Funding Sources

This work was supported by a National Health and Medical Research Council of Australia (NHMRC) project grant awarded to NES (APP1100164).

## ACKNOWLEDGEMENTS

We thank the Melbourne Mass Spectrometry and Proteomics Facility of The Bio21 Molecular Science and Biotechnology Institute for access to MS instrumentation and Byonic. We would like to thank Christine Szymanski and Justin Duma for the kind gift of *Campylobacter fetus fetus* NCTC 10842 lysates. We thank Ben Parker for feedback on the manuscript. We would also like to thank Morten Thaysen-Andersen and Daniel Kolarich for their fruitful discussions and highlighting that ZIC-HILIC enrichment was likely missing numerous Burkholderia glycopeptides. These conversations were instrumental in leading to this work.

## ABBREVIATIONS

(HILIC): Hydrophilic Interaction Liquid Chromatography
(MS): Mass spectrometry
(CV): Compensation voltage
(ZIC-HILIC): Zwitterionic hydrophilic interaction liquid chromatography
(Gal): galactose
(Glc): Glucose
(GalNAc): N-Acetylgalactosamine
(GlcNAc): N-Acetylglucosamine
(Suc): Succinyl
(diNAcBac): 2,4-diacetamido-2,4,6 trideoxyglucopyranose
(HexNAc): N-acetylhexoseamine
(Hex): Hexose
(IMS): Ion Mobility Spectrometry
(FAIMS): Field Asymmetric Waveform IMS
(LB): Luria Bertani
(PBS): Phosphate-buffered saline
(SDS): Sodium dodecyl sulfate
(NCE): Normalized collisional energy
(Tris): Tris(hydroxymethyl)aminomethane
(TFA): Trifluoroacetic acid
(DTT): Dithiothreitol
(HCD): Higher-energy collision dissociation
(CID): Collision-induced dissociation
(EThcD): Electron-transfer/higher-energy collision dissociation
(AGC): Automatic Gain Control
(DMSO): Dimethyl sulfoxide
(iBAQ): intensity-based absolute quantification
(PSMs): Peptide spectrum matches

## Notes

### Competing Interest Statement

The authors have declared no competing interest.

