## Supplementary Document (Supplementary figure 1 to 5) for "What are we missing by using hydrophilic enrichment? Improving bacterial glycoproteome coverage using total proteome and FAIMS analysis"

**Running title:** identifying HILIC inaccessible glycopeptides within bacterial glycoproteomes

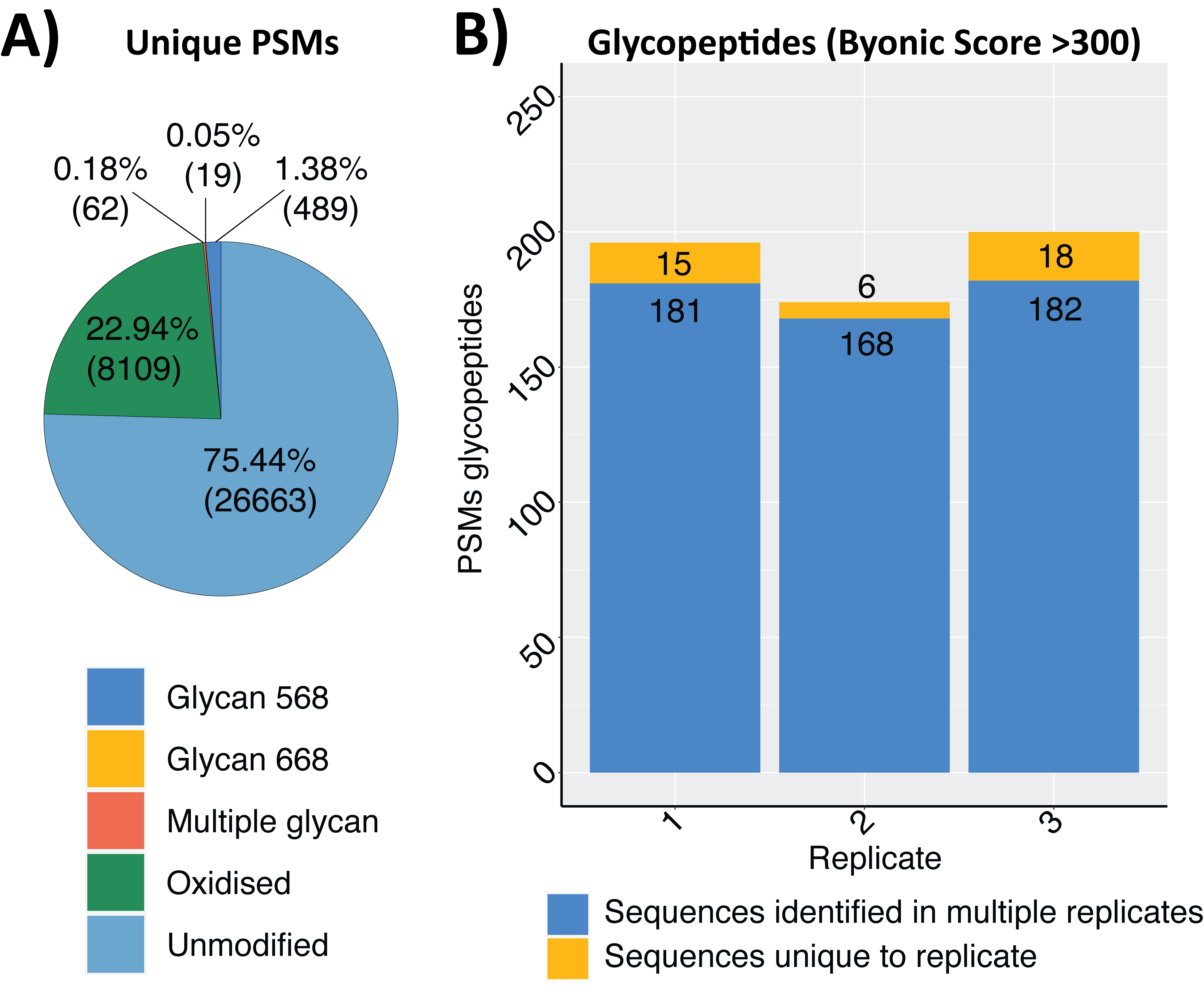

**Supplementary Figure 1: Identification of glycopeptides within *B. cenocepacia* J2315 whole proteome digests.** A) Pie charts showing all unique PSMs identified with scores above 300 B) Number of glycopeptides PSMs (Byonic Score >300) identified within each unenriched sample.

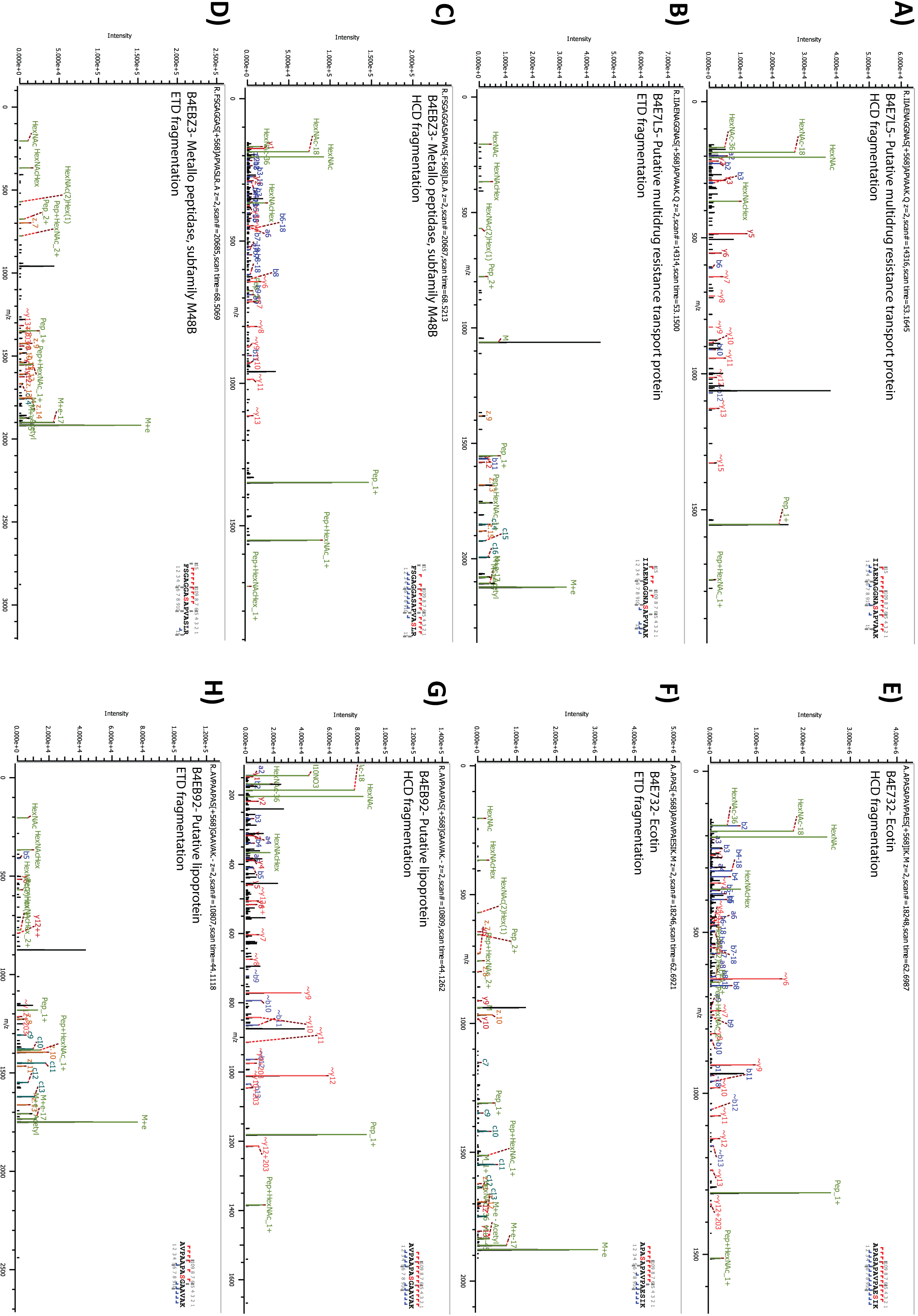

**Supplementary Figure 2: Manual assessment of glycopeptides identified with *B. cenocepacia* J2315 whole proteome digests.** HCD and ETD fragmentation of novel glycoproteins A-B) B4E7L5; C-D) B4EBZ3; E-F) B4E732 and G-H) B4EB92.

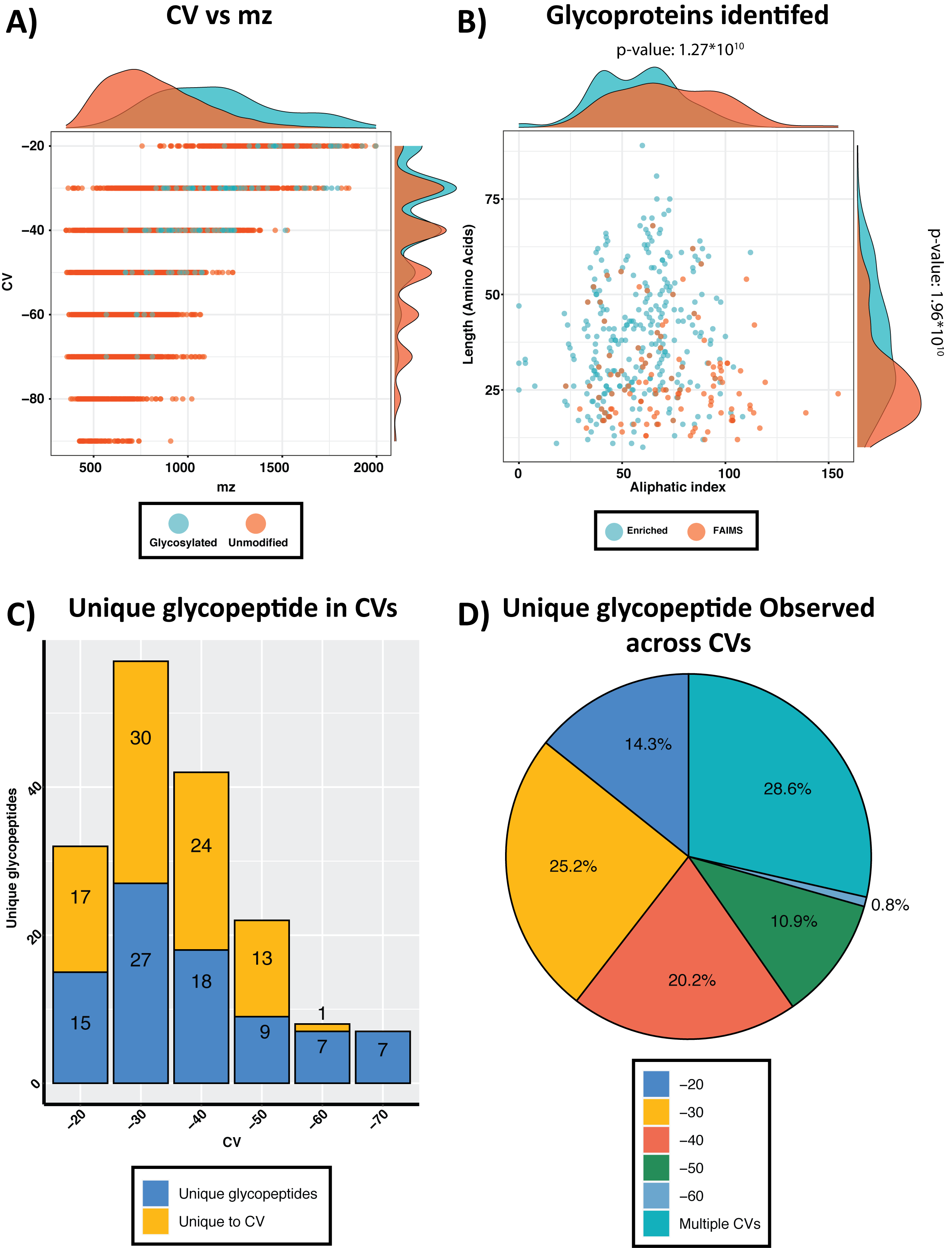

**Supplementary Figure 3: Analysis of *B. cenocepacia* glycopeptide identified within static FAIMS fractions.** A) Scatter plot of CV vs m/z of glycopeptide identified. B) Peptide properties of glycopeptide identified across static FAIMS CVs vs HILIC enriched glycopeptides

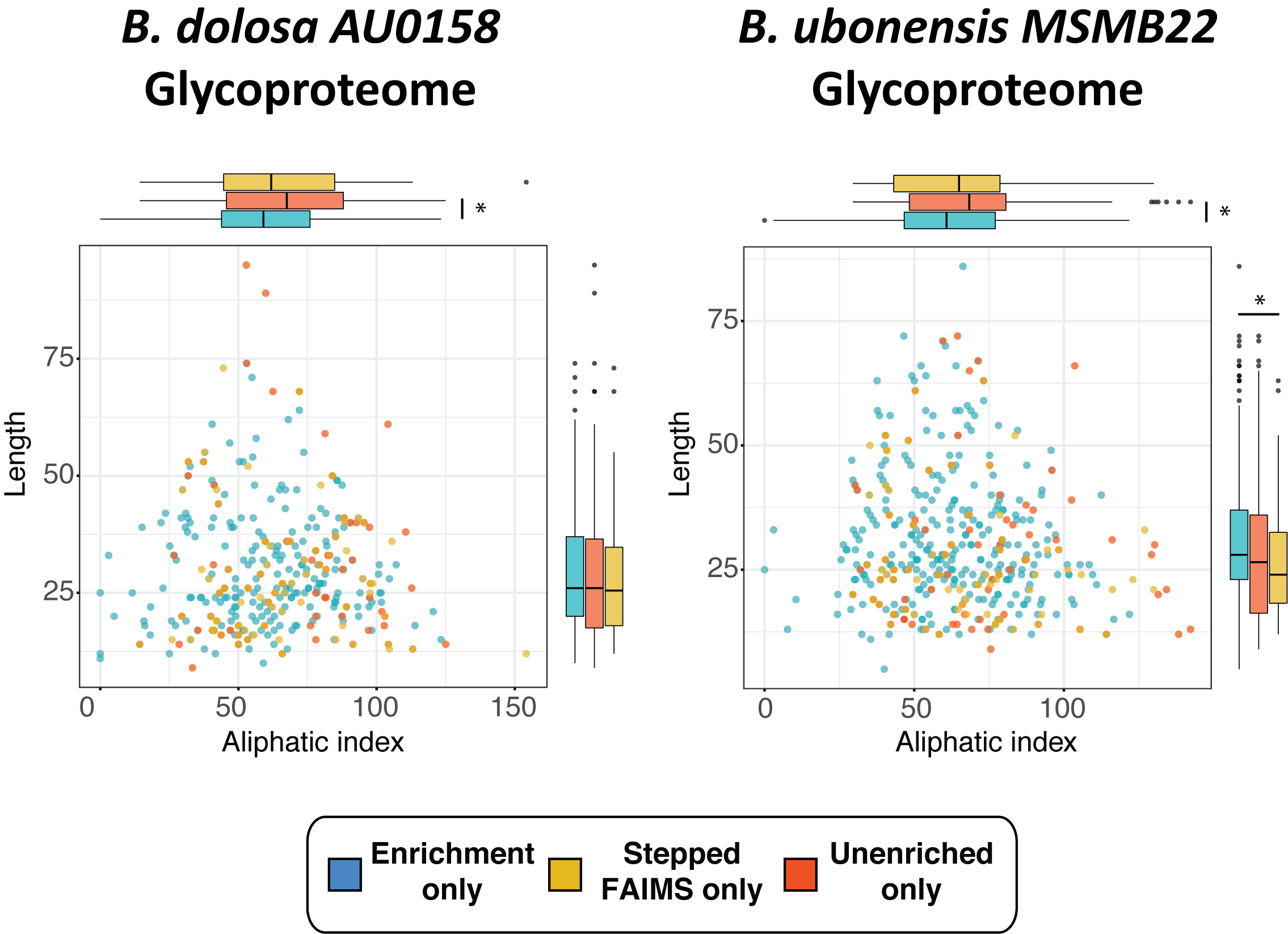

**Supplementary Figure 4: Comparisons of glycopeptide length vs aliphaticity across the Burkholderia species *B. dolosa and B. ubonensis***. Within strains examined difference in aliphaticity properties or length are observed from glycopeptides identified within stepped FAIMS and unenriched samples compared to ZIC-HILIC enrichments. Statistically significant differences in the populations are denoted with *.

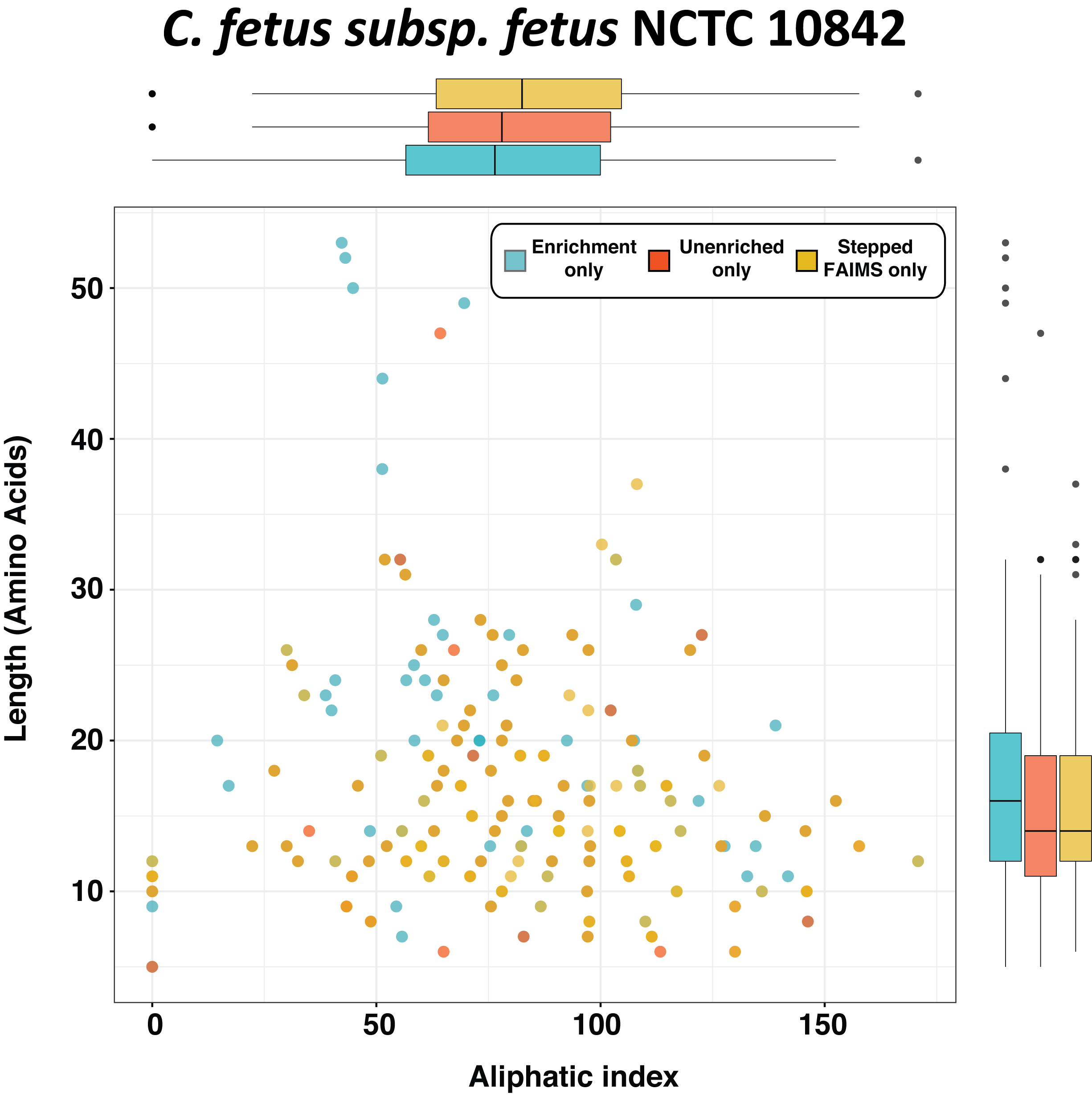

**Supplementary Figure 5: Comparisons of glycopeptide length vs aliphaticity within *C. fetus subsp. fetus glycopeptide*.** No differences in the aliphatic properties or length are observed from glycopeptides identified within stepped FAIMS and unenriched samples compared to ZIC-HILIC enrichments.
